# Inversion of a topological domain leads to restricted changes in its gene expression and affects inter-domain communication

**DOI:** 10.1101/2022.01.24.477495

**Authors:** Rafael Galupa, Christel Picard, Nicolas Servant, Elphège Pierre Nora, Yinxiu Zhan, Joke van Bemmel, Fatima El Marjou, Colin Johanneau, Maud Borensztein, Katia Ancelin, Luca Giorgetti, Edith Heard

**Affiliations:** Mammalian Developmental Epigenetics Group, Genetics and Developmental Biology Unit, Institut Curie, PSL Research University, CNRS UMR3215, INSERM U934, Paris, France; Bioinformatics, Biostatistics, Epidemiology and Computational Systems Unit, Institut Curie, PSL Research University, INSERM U900, Paris, France; MINES ParisTech, PSL Research University, CBIO-Centre for Computational Biology, Paris, France; Friedrich Miescher Institute for Biomedical Research, Basel, Switzerland; University of Basel, Switzerland; Transgenesis Facility, Institut Curie, Paris, France; Collège de France, Paris, France

**Author notes:** Correspondence to and; current address: European Molecular Biology Laboratory, Heidelberg, Germany. Equal contribution (Rafael Galupa, Christel Picard). Cardiovascular Research Institute, University of California San Francisco, San Francisco, CA, USA; Department of Biochemistry and Biophysics, University of California San Francisco, San Francisco, CA, USA.

## Abstract

The interplay between the topological organization of the genome and the regulation of gene expression remains unclear. Depletion of molecular factors underlying genome topology, such as CTCF and cohesin, leads to modest alterations in gene expression, while genomic rearrangements involving boundaries of topologically associating domains (TADs) disrupt normal gene expression and can lead to pathological phenotypes. Here we inverted an almost entire TAD (245kb out of 300kb) within the *X-inactivation centre* (*Xic*), leaving its boundaries intact. This led to a significant rearrangement of topological contacts within the TAD, mostly in accordance to the orientation of underlying CTCF binding sites but suggesting heterogeneity in the “contact” potential of different CTCF sites. The inversion also led to increased contact insulation with the neighbouring TAD. Expression of most genes within the inverted TAD remained unaffected in mouse embryonic stem cells and during differentiation. Interestingly, expression in the neighbouring TAD of the noncoding transcript *Xist*, which controls X-chromosome inactivation, was ectopically upregulated. The same inversion in mouse embryos led to a bias in *Xist* expression, but X-inactivation choice ratios did not significantly deviate from wild type. Smaller deletions and inversions of specific clusters of CTCF sites within the TAD led to similar results: rearrangement of contacts, limited changes in local gene expression but significant changes in *Xist* expression. Our study suggests that the wiring of regulatory interactions within a TAD can influence the expression of genes in neighbouring TADs, highlighting the existence of mechanisms for inter-TAD communication.

## Introduction

The three-dimensional folding of the genome has been increasingly recognised as an essential component for our understanding of gene regulation (Dekker and Mirny, 2016; McCord et al., 2020). Chromosome conformation capture techniques (Denker and de Laat, 2016) have unravelled a complex hierarchy of structural layers that organise mammalian chromosomes, composed of domains of high frequency contacts (Zhan et al., 2017). At the sub-megabase level, these domains are generally designated topologically associating domains (TADs) (Dixon et al., 2012; Nora et al., 2012) and are well conserved across species and invariant across cell types (Dekker and Heard, 2015). The dynamics of the formation and maintenance of TADs and their boundaries during development and each cell cycle remain under investigation (Szabo et al., 2019) but seem to depend on the interplay between the architectural proteins cohesin and the zinc finger protein CTCF (Fudenberg et al., 2016; Haarhuis et al., 2017; Nora et al., 2017; Rao et al., 2017; Sanborn et al., 2015; Schwarzer et al., 2017; Wutz et al., 2017). Enriched at boundaries between TADs (Dixon et al., 2012; Phillips-Cremins et al., 2013), CTCF is required for chromatin loops observed between CTCF sites and for the organisation and insulation of most TADs (Nora et al., 2017). Remarkably, CTCF-mediated contacts mostly occur between CTCF sites for which the CTCF motifs lie in a convergent orientation (Rao et al., 2014a; Tang et al., 2015a), and the contacts depend on the orientation of the motifs: altering the orientation of a CTCF site can disrupt a loop and lead to the formation of new ones (de Wit et al., 2015; Guo et al., 2015; Sanborn et al., 2015).

TADs are thought to instruct gene regulatory landscapes, allowing promoters and their regulatory elements to meet often and lead to a more efficient transcriptional output (Symmons et al., 2016). Accordingly, TADs represent the folding scale at which promoter-enhancer interactions and gene co-regulation are maximised (Zhan et al., 2017). The communication between promoters and enhancers is generally assumed to rely on chromatin looping, and long-range contacts within TADs can be quite dynamic during processes that involve rewiring of the regulatory networks, such as differentiation (Dixon et al., 2015). However, the interplay between such topological organization and the regulation of gene expression remains unclear. Loss of TADs upon depletion of CTCF or cohesin leads to relatively small effects on gene expression (Nora et al., 2017; Rao et al., 2017; Schwarzer et al., 2017; Wutz et al., 2017), and genomic rearrangements involving mammalian TADs and their boundaries can have either very modest effects (Amândio et al., 2020; Despang et al., 2019; Paliou et al., 2019; Rodríguez-Carballo et al., 2017; Williamson et al., 2019) or disrupt normal gene expression and underlie pathological phenotypes (Flavahan et al., 2016; Franke et al., 2016; Hnisz et al., 2016; Lupiáñez et al., 2015).

Here we set out to explore the relationship between TAD organisation and transcriptional regulation at a critical developmental regulatory landscape, the mouse *X-inactivation centre* (*Xic*). The *Xic* is the master regulator for the initiation of X-chromosome inactivation in female placental mammals (Augui et al., 2011a; Rastan and Brown, 1990), harbouring the noncoding RNA *Xist* locus and the regulatory elements necessary for its female-specific developmental control. *Xist* is repressed in mouse embryonic stem cells (mESCs) and becomes upregulated from one of the two X-chromosomes in females upon exit of the pluripotent state, leading to random X-inactivation. This upregulation depends on *Xist* cis-regulatory landscape (Heard et al., 1999), the full extent of which is still undefined – it is, however, partitioned in at least two TADs, with the *Xist* locus lying close to the boundary between them (**Fig. 1A**) (Nora et al., 2012). The TAD in which the *Xist* promoter is included (here referred to as Xist-TAD) contains some of *Xist* positive regulators (Augui et al., 2007; Barakat et al., 2011, 2014; Furlan et al., 2018; Gontan et al., 2012; Jonkers et al., 2009; Tian et al., 2010), while the adjacent TAD (here referred to as Tsix-TAD) contains the promoter of *Tsix*, the antisense transcription unit to *Xist* that blocks its upregulation (Lee and Lu, 1999; Luikenhuis et al., 2001; Stavropoulos et al., 2001) as well as other elements that act as a cis-repressors of *Xist* (such as *Linx* and *Xite*, see more below).

**Figure 1.**
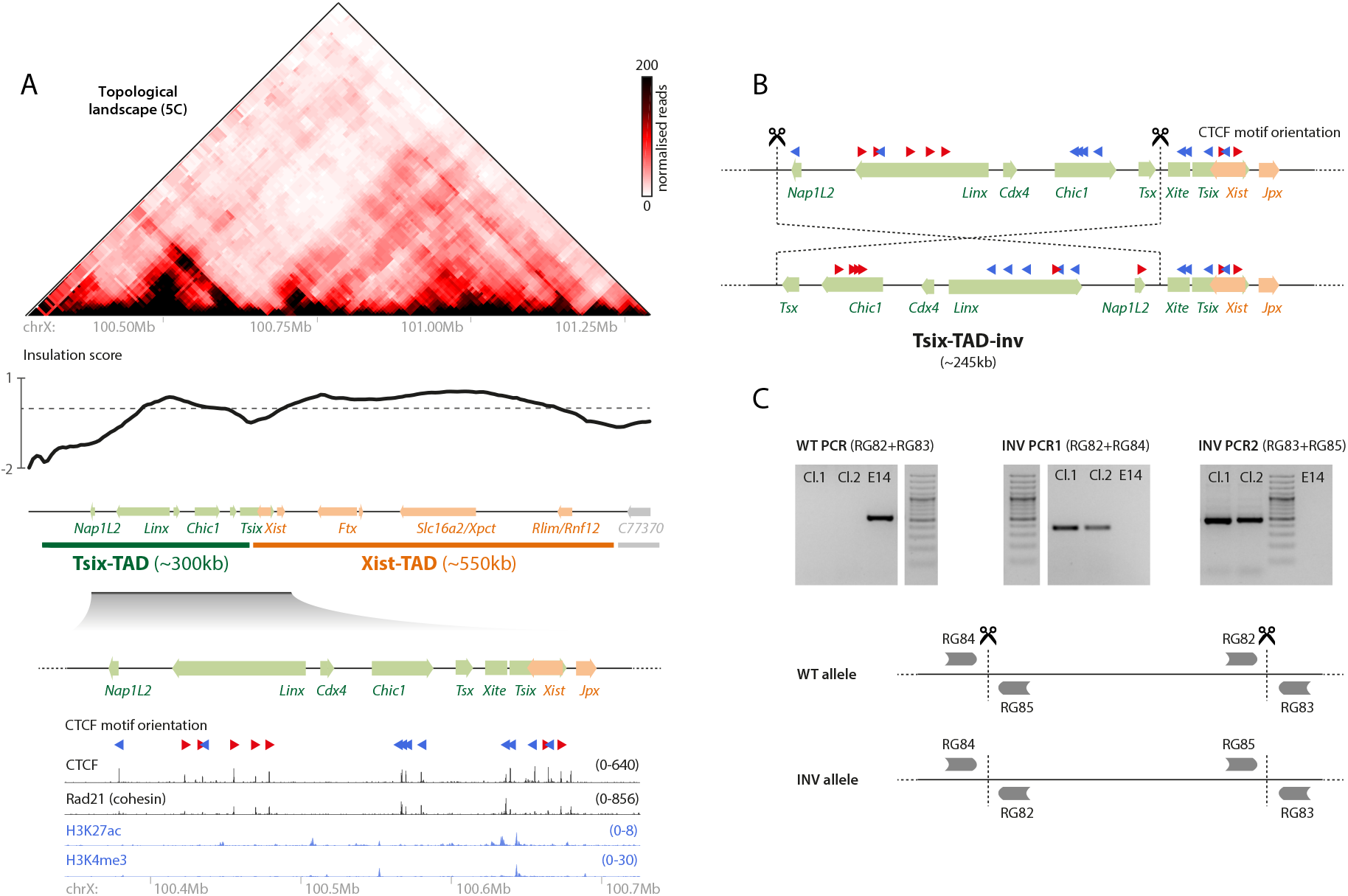
Strategy for inverting the nearly entire Tsix-TAD. (*A*) Topological organisation of the *Xic* (top) and chromatin ChIP-seq profiles (bottom; see Methods for sources); the *Xist*/*Tsix* locus lies at the boundary between two TADs. (*B*) Targeting strategy for inverting the ∼245kb region comprising most of the Tsix-TAD, except *Tsix* and its known regulator *Xite*, and leaving the boundaries intact. (*C*) PCR strategy (bottom) and gel results (top) for detecting the inversion events. E14 is the wild type, parental cell line. Cl.1 and Cl.2 are the two clones that were generated and analysed throughout the study.

To investigate how the topological organisation of the Tsix-TAD impacts the regulation of genes therein, and in the neighbouring Xist-TAD, we generated a mutant allele in mESCs and in mice with an inversion of almost the entire Tsix-TAD (245kb out of 300kb). We found that rewiring the Tsix-TAD structural landscape led to the formation of new chromatin contacts within the TAD, generally following the folding principles determined by the orientations of CTCF motifs. These topological alterations were accompanied by changes in gene expression of two out of seven genes within the TAD in differentiating mESCs. Interestingly, we found that the expression of *Xist*, in the neighbouring TAD, was ectopically upregulated, suggesting that inter-TAD communication was affected

## Results

### Generating a genomic inversion encompassing the Tsix-TAD (245kb-INV)

The Tsix-TAD harbours three hotspots of physical contacts (**Fig. 1A**), involving three different loci: (1) the *Xite* element, a proximal enhancer of *Tsix* (Ogawa and Lee, 2003) also involved in the position and insulation of the boundary between the Tsix-and Xist-TADs (van Bemmel et al., 2019); (2) the noncoding *Linx* locus, which harbours two cis-regulatory elements involved in controlling *Xist* expression (Galupa et al., 2020); and (3) *Chic1*, previously implicated in the maintenance of the organisation of the Tsix-TAD (Giorgetti et al., 2014). Each of these loci harbours a set of CTCF sites involved in mediating the observed physical contacts, and within each locus, most CTCF motifs present the same orientation (**Fig. 1A**). Sites within *Linx* are “convergently oriented” towards those within *Chic1* or *Xite*, the preferred orientation to form chromatin loops (Rao et al., 2014a; Tang et al., 2015b). Contacts between *Chic1* and *Xite* (the CTCF motifs of which occur in a “tandem”) are also observed (**Fig. 1A**). The contacts between these three loci might occur in pairwise fashion and/or simultaneously; physical modelling suggests that all conformations are possible (Giorgetti et al., 2014) and deletions of the CTCF binding sites in either *Xite* (van Bemmel et al., 2019) or *Linx* (Galupa et al., 2020) show that contacts between the two remaining loci still occur.

We wondered whether this complex topological organization might be critical to ensure correct communication between the surrounding cis-regulatory elements (such as those within *Xite* and *Linx*) and their targets, and therefore ensure appropriate gene expression of *Tsix* and *Xist* and correct patterns of X-inactivation. Using a CRISPR/Cas9 editing approach in male mESCs, which carry a single X chromosome, we targeted a ∼245kb region encompassing all loci within the Tsix-TAD, including the CTCF clusters within *<u>Linx</u>* and *Chic1*, but excluding *Xite* and *Tsix* (**Fig. 1B**). We decided not to include *Xite* in the inversion because (i) *Xite* is already known to influence *Xist* expression (via *Tsix*), and (ii) if *Xite* was inverted along with the rest of the TAD, the relative CTCF orientations between *Xite, Linx* and *Chic1* would not have changed. The targeted region does not involve either of the two boundaries of the TAD. We successfully generated two clones harbouring an inversion allele (245kb-INV) (**Fig. 1C**). This genomic inversion swaps the orientations of all CTCF motifs therein relative to those outside of the inverted region, in particular for *Linx* and *Chic1* (**Fig. 1B**), and is therefore expected to lead to the formation of new contacts within the TAD.

### 245kb-INV leads to rearrangement of contacts within the TAD and increased insulation with neighbouring TAD

To assess the topological organisation of the 245kb-INV allele, we performed carbon-copy chromosome conformation capture (5C) on the *Xic* (Dostie et al., 2006; Nora et al., 2012) for mutant and control mESCs (**Fig. 2A**). 5C analysis revealed that three hotspots of contacts can still be observed in the Tsix-TAD on the 245kb-INV allele (**Fig. 2B;** please note that the 5C map is shown after “correction” of the new genomic sequences in the inverted allele). These involve the same three loci as in control cells: in its new position, the *Chic1* CTCF cluster is able to establish contacts with *Linx* and with *Xite* (**Fig. 2B**) and *Linx* and *Xite*, with CTCF sites in “tandem” orientation in the 245kb-INV allele, also interact together (like *Chic1* and *Xite* do in control cells) (**Fig. 2B**). Inverting the *Linx* and *Chic1* CTCF clusters simultaneously seems therefore to lead to new but similar hotspots of physical contacts within the Tsix-TAD compared to control. This could have been expected given that the overall distribution and orientation of CTCF sites within the TAD remains similar between the wild type and the inverted alleles (**Fig. 1B**). In other words, the *Chic1* CTCF cluster on the inverted allele occupies an equivalent position to *Linx* on the wild type allele, and vice-versa. Therefore, the 245kb inversion can lead to the formation of similar loops within the Tsix-TAD compared to wild type – yet, involving different cis-regulatory elements.

**Figure 2.**
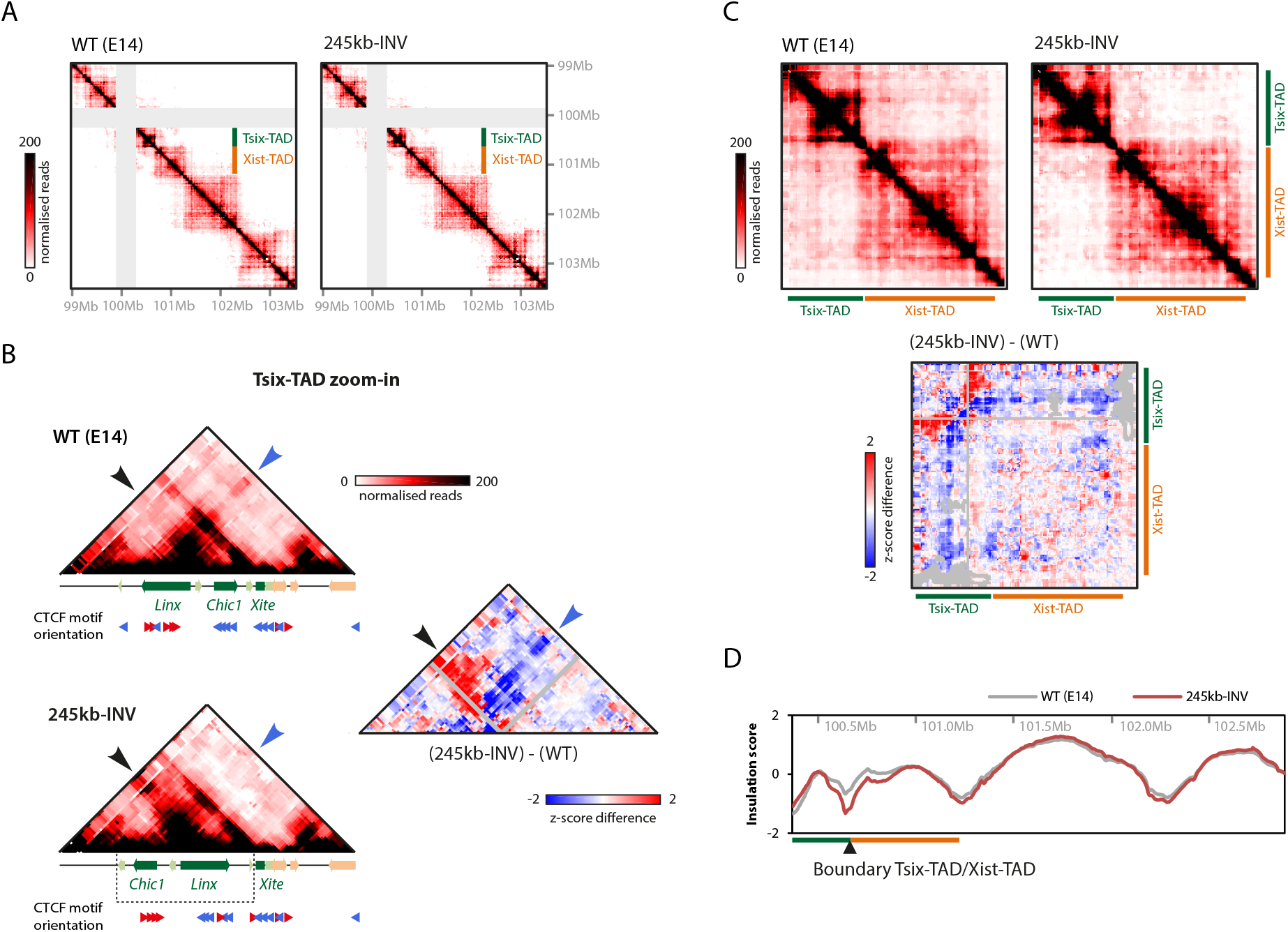
Rearrangement of contacts within the TAD and increased insulation with neighbouring TAD upon 245kb intra-TAD inversion. (*A*) 5C profiles of wildtype (WT; two replicates pooled) and 245kb-INV mutant (two clones pooled) mESCs. Mutant map is corrected for inversion and grey pixels represent filtered contacts (see Methods). (*B*) (*C*) Zoom-ins of the maps in (A) and 5C differential maps, representing the subtraction of Z-scores calculated for wildtype and 245kb-INV mutant maps separately. Grey pixels represent filtered contacts. (*D*) Insulation scores across the *Xic* TADs and downstream TADs based on 5C profiles for wildtype and 245kb-INV mutant mESCs. The “valleys” represent TAD boundaries.

Nevertheless, we also noticed some significant differences in the topology of the “inverted” Tsix-TAD. Increased contacts could be observed upstream of the inverted region, corresponding to contacts stemming from the *Linx* CTCF cluster in its new position (**Fig. 2B**, bottom, black arrow, red region in the differential map; this region shows no particular chromatin signatures, such as CTCF binding or active chromatin marks). This suggests a different “strength” for the *Linx* and *Chic1* CTCF clusters: in the inverted allele, the *Linx* CTCF cluster strongly interacts with regions upstream of *Chic1* (**Fig. 2B**, bottom, black arrow), while in the wild type configuration, the *Chic1* CTCF cluster does not form such strong contacts with regions upstream of *Linx* (**Fig. 2B**, top, black arrow). Conversely, we also observed a localised, strong reduction in contacts (**Fig. 2B**, differential map, blue arrow) associated with the switch in positions between *Linx* and *Chic1*: the *Linx* CTCF cluster at its original position is able to form long-range contacts beyond *Chic1* and *Xite*, with elements within the Xist-TAD (**Fig. 2B**, top, blue arrow). These contacts are lost (or strongly reduced) in the 245kb-INV cells (**Fig. 2B**, bottom and differential map, blue arrows), indicating that the *Chic1* CTCF cluster does not establish long-range contacts with the Xist-TAD when placed in the *Linx* CTCF cluster position. This loss of contacts across the boundary actually extends along the whole Xist-TAD (**Fig. 2C**). Again, this suggests a stronger potential for the *Linx* CTCF cluster to form contacts, compared to the *Chic1* CTCF cluster.

We also evaluated to which extent these topological changes had an impact in the overall insulation of the TADs. Insulation score analysis (see Methods) revealed a clear gain of insulation across the boundary between the Tsix-TAD and the Xist-TAD (**Fig. 2D**; lower insulation scores are reflective of increased insulation). The loss of Linx-mediated contacts across the boundary probably accounts for this increased insulation between the TADs, at least partially. In summary, the 245kb inversion repositions CTCF clusters within the Tsix-TAD, leading to a reconfiguration of specific intra- and inter-TAD contacts accompanied by stronger insulation with the neighbouring Xist-TAD.

### 245kb-INV leads to gene expression changes within the Tsix-TAD and across the boundary

We next set out to understand if similar interaction patterns, but different wiring of sequences within the Tsix-TAD, led to any transcriptional changes. To this end, we profiled transcript levels across the *Xic* using digital gene expression analysis (NanoString) (Geiss et al., 2008) in control and mutant cells in the pluripotent state (d0) and during early differentiation (d0.5-d2.5) (**Fig. 3A**). Expression of most genes within the Tsix-TAD and the Xist-TADs was unaffected in 245kb-INV cells (**Fig. 3B**), including that of the three loci involved in the topological alterations, *Linx, Chic1* and *Xite*. This suggests no or limited effect of the structural alterations on the transcriptional regulation of these loci. Expression of *Tsix* was significantly reduced in mutant cells in the pluripotent state (d0) (**Fig. 3B**) but such effect did not persist consistently during differentiation. The deletion of the same region that we inverted here also led to downregulation of *Tsix* in mESC (Galupa et al., 2020); together with the current data, this suggests that the region contains important sequences for *Tsix* regulation and that this regulation depends on the orientation of the region as a whole, and might depend on the orientation of individual regulatory sequences.

**Figure 3.**
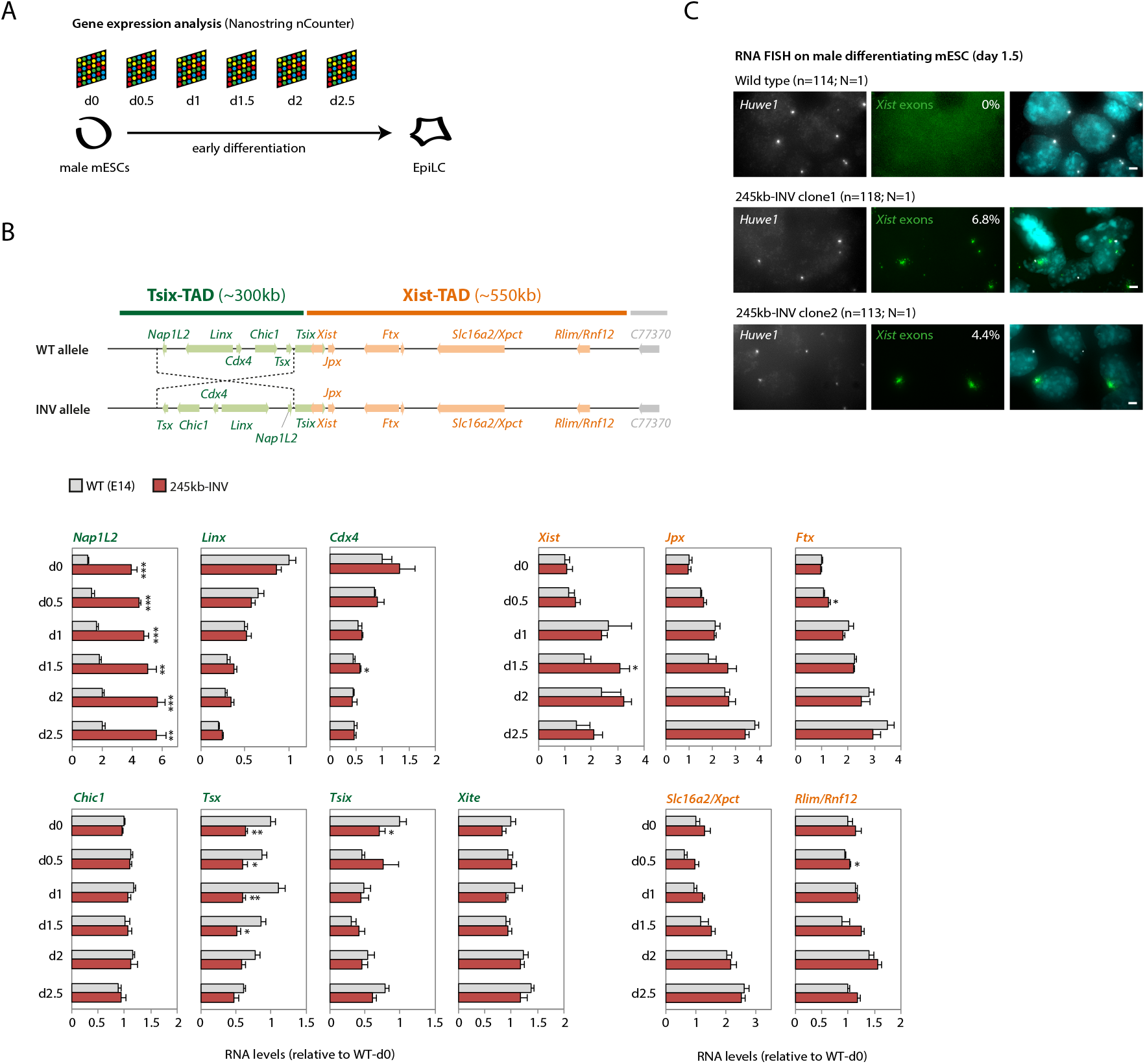
Inversion leads to transcriptional changes of specific genes within the TAD and of *Xist* across the TAD boundary. (*A*) Schematic representation of mESC to EpiLSC differentiation and time points analysed by Nanostring nCounter (see Methods). (*B*) Gene expression analysis during differentiation. Data is normalised to wt-d0 for each gene, and represents the average of two biological replicates (wild type) or of two independent clones (mutant). Statistical analysis: two-tailed paired t-test (* p<0.05; ** p<0.01; *** p<0.001). (*C*) RNA FISH for Huwe1 (X-linked gene outside of the *Xic*) and Xist (exonic probe) on mESCs differentiated to d1.5. Percentage of cells with Xist RNA accumulation is indicated and represents an average from two independent experiments. Scale bar: 2 1m.

We did notice however consistent changes during differentiation in mutant cells for two genes within the Tsix-TAD: *Nap1L2*, which was significantly upregulated at all time points (**Fig. 3B**) and *Tsx*, which was significantly downregulated (**Fig. 3B**). Interestingly, both genes lie at the extremities of the inverted region, and switch their relative positions in the TAD between the wild type and mutant configurations. It is likely that changes in their gene expression are associated with altered proximity to the *Xite* enhancer element. Since deletion of *Xite* leads to downregulation of *Tsx* (van Bemmel et al., 2019), moving *Tsx* away from *Xite* on the 245kb-INV allele could lead to its observed downregulation. Conversely, increased linear proximity of *Nap1L2* to *Xite* could possibly underlie *Nap1L2* upregulation. Changes in interaction frequencies between *Xite* and these two elements in the 245kb-INV allele do support this hypothesis, as they reflect the changes in their genomic distances (increased for *Xite-Nap1L2* and decreased for *Xite-Tsx*, compared with control, **Fig. S1**).

We also observed changes in expression of *Xist*, the long noncoding RNA locus that is regulated by the *Xic* to trigger the initiation of X-chromosome inactivation. Normally very low in male cells, *Xist* expression was slightly upregulated in the mutants at later differentiation time points (∼2-fold at d1.5; **Fig. 3B**). In female cells undergoing X-inactivation, upregulation of *Xist* is accompanied by local accumulation of its RNA in *cis*, forming a so-called “Xist cloud”, which can readily be detected by RNA FISH (Augui et al., 2011b). RNA FISH revealed the formation of Xist clouds in ∼4-7% of mutant male cells upon differentiation, which was never observed in wild type male cells (**Fig. 3C**). Thus, the inversion of 245kb within the Tsix-TAD leads to ectopic expression of *Xist*, the promoter of which is located in the neighbouring TAD.

### Female embryos with 245kb-INV allele show higher Xist allelic imbalance

Given the impact of the 245kb inversion on *Xist* expression in male cells, we decided to investigate whether this was also the case in female embryos at post-implantation stages, when random X-inactivation is known to have already occurred (Rastan, 1982). To this end, we generated an equivalent 245kb-INV allele in mice (see Methods) and collected post-implantation heterozygous embryos. These embryos were derived from crosses between polymorphic mouse strains (**Fig. 4A, 4D**), which allows us to distinguish the allelic origin of transcripts. Analysis of RNA allelic ratios for *Atp7a*, an X-linked gene, revealed no preferential gene silencing for one or the other allele (**Fig. 4B, 4E**).

**Figure 4.**
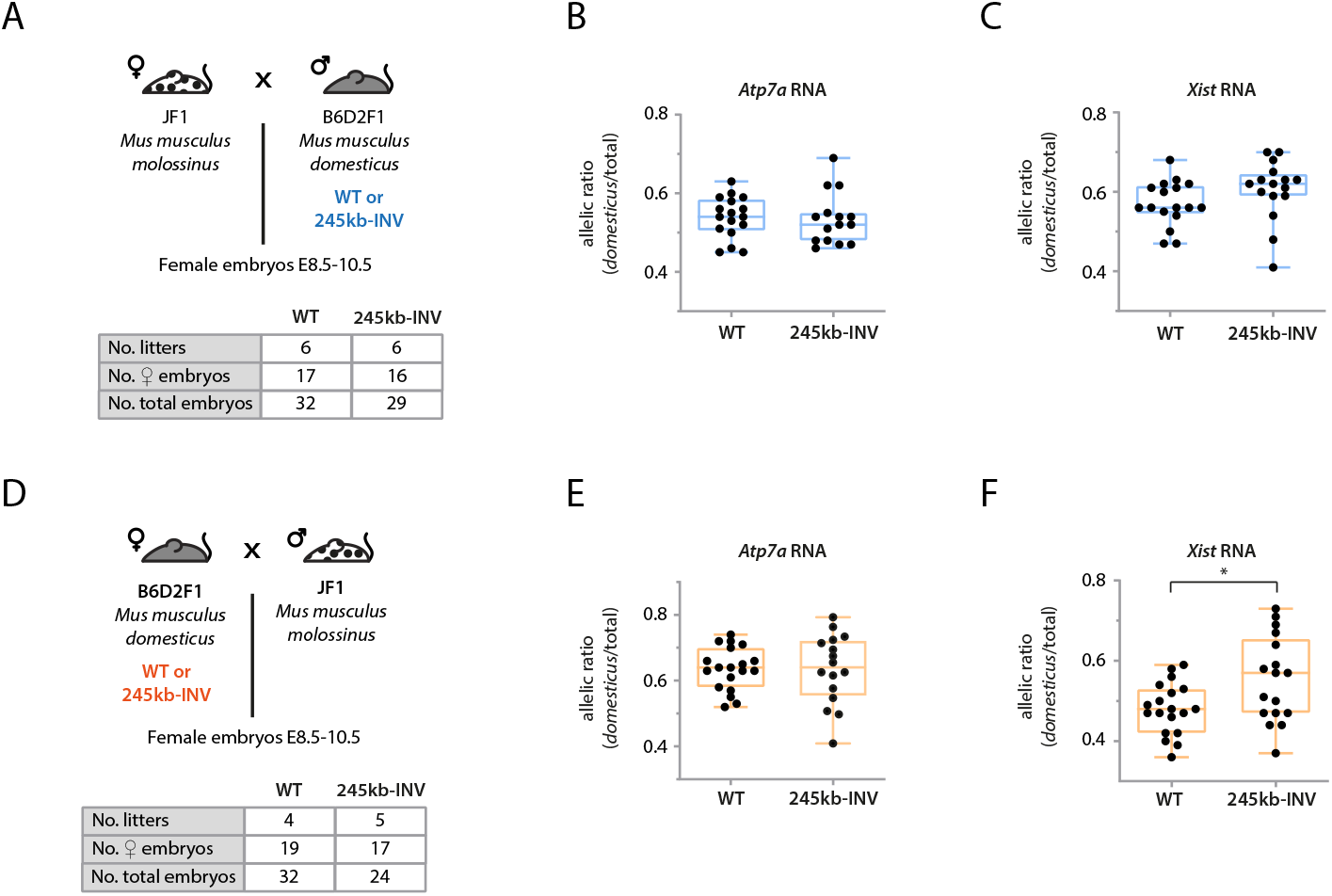
Female embryos with 245kb-INV allele show a bias in Xist expression. (*A, D*) Crosses used for analysis of RNA allelic ratios in female hybrid embryos inheriting the *M. musculus domesticus* allele paternally (*A*) or maternally (*D*). Tables summarise number of embryos collected. (*B, C, E, F*) RNA allelic ratios for the X-linked gene *Atp7a* (*B, E*) and *Xist* (*C, F*). Each black dot corresponds to a single female embryo. Statistical analysis was performed using Mann-Whitney test (see Methods).

However, analysis of *Xist* RNA allelic ratios between mutant and control embryos showed slightly higher *Xist* allelic ratios in the mutant embryos, whether the mutant allele was inherited paternally (**Fig. 4C**) or maternally (**Fig. 4F**); this increase was statistically significant for maternal transmission (p<0.05). These results are consistent with the upregulation of *Xist* that we observed in mutant cells (**Fig. 3B-C**). Thus, the 245kb inversion leads to higher *Xist* levels in *cis* but this does not result in skewed patterns of X-inactivation. Of note, litter size seems to be reduced upon maternal transmission of the 245kb-INV allele, with a skewed sex ratio (71% females in 245kb-INV, 59% in control) suggesting that the inversion may have more phenotypic consequences.

### Mutating clusters of CTCF sites within Linx and Chic1 lead to changes in Xist expression

To further explore the link between the topological organization of the Tsix-TAD and *Xist* regulation, we decided to generate alleles with deletions and/or inversions of the clusters of CTCF sites within *Linx* and within *Chic1*. We have previously deleted a large intronic interval containing three *Linx* CTCF sites, in male ESCs (∼51 kb) and in mice (∼25 kb) (Galupa et al., 2020), which led to some alterations in the topological organization of the two *Xic* TADs but no changes in *Xist* expression in female embryos. We decided to test the impact of inversions of exactly the same regions, in male mESCs (Linx-51kb-INV) and in mice (Linx-25kb-INV) (**Fig. 5A**). 5C analysis of the Linx-51kb-INV allele revealed higher frequency of contacts between the now inverted *Linx* locus and regions immediately upstream (**Fig. 5B**, black arrowhead), and lower frequency of contacts between (inverted) *Linx* and *Chic1, Xite* and elements within the Xist-TAD (**Fig. 5B-C**, blue arrowhead), in agreement with the change of orientation of the three *Linx* CTCF sites. These results are reminiscent of what we observed for the 245kb-INV allele (**Fig. 2B-C**), and they support the hypothesis that loss of contacts with the Xist-TAD in the 245kb-INV allele is associated to inversion of the CTCF sites within *Linx*. Consistently, analysis of insulation scores across the TADs revealed a gain of insulation across the boundary between the Tsix-TAD and the Xist-TAD (**Fig. 5D**), though less pronounced than what we observed for 245kb-INV allele (**Fig. 2D**). We next analysed gene expression across the *Xic* for the Linx-51kb-INV mESC in the pluripotent state (d0) and during early differentiation (d0.5-d2.5); expression of *Linx* was significantly downregulated at some time points (**Fig. 5E**) but no changes were observed for *Xist* or *Tsix* (**Fig. 5E**) nor for any other locus across the *Xic* (data not shown). However, when we analysed *Xist* expression in female embryos carrying an heterozygous Linx-25kb-INV allele, we observed significantly higher expression of *Xist* for the inverted allele, for both paternal and maternal transmission (**Fig. 5F-G**), and also corresponding decrease in expression of the X-linked gene *Atp7a* (**Fig. 5F-G**), suggestive of skewed XCI compared to control. Overall, the inversion of the *Linx* CTCF cluster leads to similar phenotypes compared to the large 245kb inversion, namely a decrease in contact frequency between *Linx* and the Xist-TADs, and a concomitant gain of insulation between them, and increased *Xist* expression in cis in female embryos.

**Figure 5.**
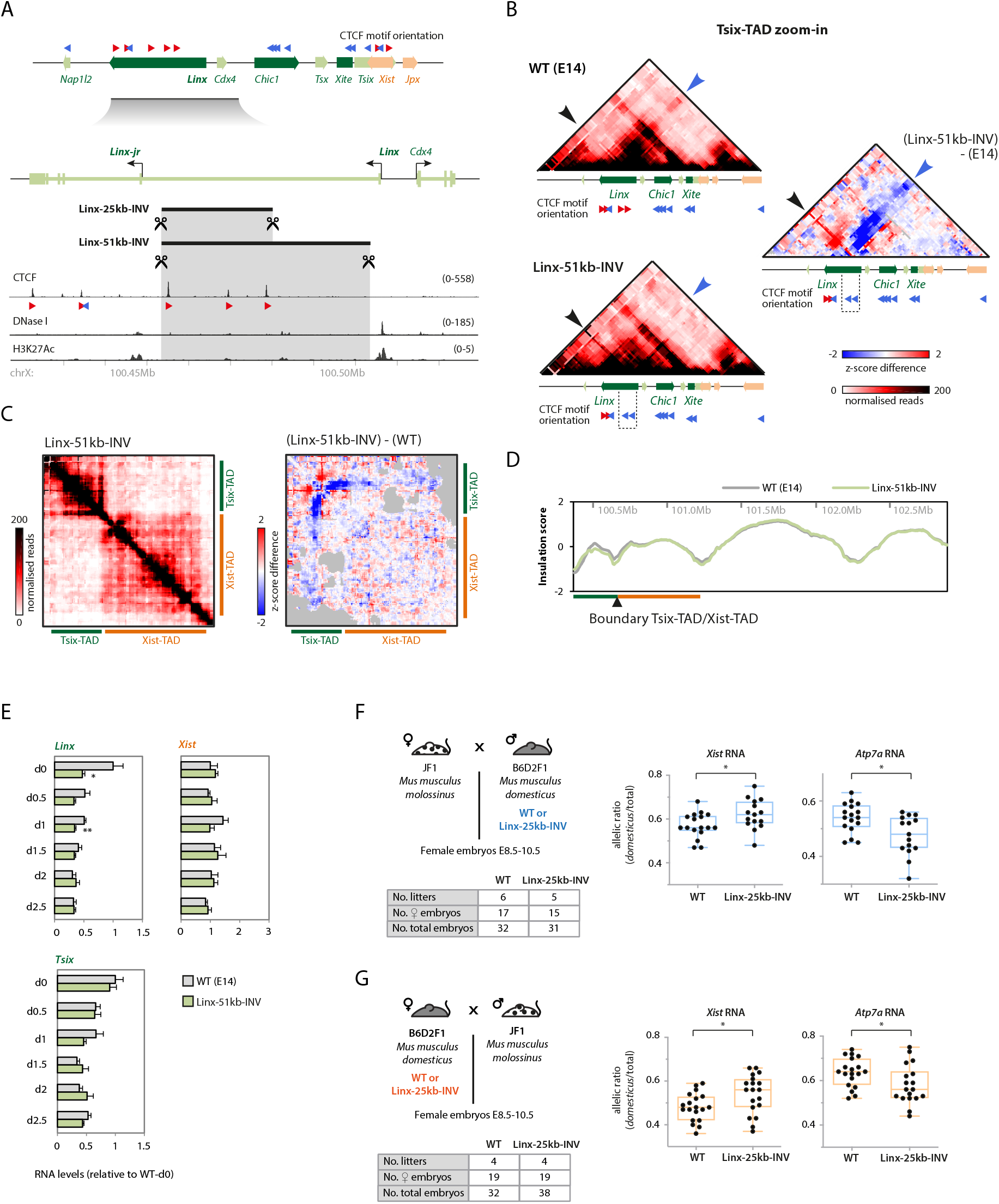
Inversion of *Linx* cluster of CTCF sites leads to Xist upregulation in *cis*. (*A*) The *Linx* locus, CTCF binding, and orientation of CTCF motifs associated with CTCF chromatin immunoprecipitation sequencing (ChIP-seq) peaks. The targeted inversions Linx-25kb-INV and Linx-51kb-INV are indicated. (*B*) 5C profiles (Tsix-TAD zoom-in) of wildtype (WT; two replicates pooled) and Linx-51kb-INV (two clones pooled) mESCs, and 5C differential map, representing the subtraction of Z-scores calculated for wildtype and Linx-51kb-INV maps. (*C*) Left, 5C profile of Linx-51kb-INV mESCs (two clones pooled); map is corrected for inversion and grey pixels represent filtered contacts (see Methods). Right, 5C differential map, representing the subtraction of Z-scores calculated for wildtype and Linx-51kb-INV maps separately. (*D*) Insulation scores across the *Xic* TADs and downstream TADs based on 5C profiles for wildtype and Linx-51kb-INV mESCs. The “valleys” represent TAD boundaries. (*E*) Gene expression analysis during differentiation. Data is normalised to wt-d0 for each gene, and represents the average of two biological replicates (wild type) or of two independent clones (mutant). Statistical analysis: two-tailed paired t-test (all nonsignificant). (*F, G*) Left, crosses used for analysis of RNA allelic ratios in female hybrid embryos inheriting the *M. musculus domesticus* allele paternally (*F*) or maternally (*G*). Tables summarise number of embryos collected. Right, RNA allelic ratios for *Xist* and the X-linked gene *Atp7a*. Each black dot corresponds to a single female embryo. Statistical analysis was performed using Mann-Whitney test (* p<0.05).

We have previously generated in male mESC a ∼4kb deletion within *Chic1* (Giorgetti et al., 2014) that encompassed two of the three CTCF binding sites present in the locus (**Fig. 6A**), but we did not study its impact on chromosome conformation nor on *Xist* expression, which we set out to do here. Differential 5C analysis between this Chic1-4kbΔ allele and wildtype revealed showed a reduction in contacts between *Chic1* and *Linx*, and also between *Chic1* and *Xite* (**Fig. 6B**), consistent with loss of the *Chic1* CTCF sites. We also noted a seemingly increase in contact frequency between *Linx* and *Xite* (**Fig. 6B, Fig. S1**), which would be consistent with a model of competition between *Chic1* and *Xite* CTCF sites to form loops with the CTCF sites within *Linx*. However, these differences in contact frequencies overall remained rather close to the “noise” levels of the 5C map. We wondered whether these effects would be more pronounced if the remaining CTCF binding site was also removed; thus, we generated, in male mESC, a larger deletion (Chic1-14kbΔ) encompassing all three CTCF sites within Chic1 (**Fig. 6A**). We observed similar contact rearrangements within the Tsix-TAD as for Chic1-4kbΔ but more pronounced (**Fig. 6C**), suggesting that it is the loss of the CTCF sites that underlies the observed topological differences. To study the impact of these deletions on gene expression across the *Xic*, we profiled transcript levels as done previously in the pluripotent state (d0) and during early differentiation. Expression of *Chic1* itself was consistently upregulated in both Chic1-4kbΔ and Chic1-14kbΔ (**Fig. 6D-E**); it is intriguing to think that this could potentially be linked to its now shorter length, as shorter genes have been associated to higher levels of expression (Castillo-Davis et al., 2002; Chiaromonte et al., 2003). We also observed higher expression of the gene upstream of *Chic1, Cdx4*: interestingly, the effects seemed to scale up with the larger deletion – in Chic1-4kbΔ mESC, there was a slight increase in *Cdx4* levels across time points but not statistically significant, while in Chic1-14kbΔ the increase was more pronounced and statistically significant for some of the time points. This effect could be connected to the removal of all CTCF sites from within the *Chic1* locus, which could potentially “shield”, or insulate, *Cdx4* from activating influences downstream of the CTCF sites. Expression of *Xist* expression was also more affected in mESC containing the larger deletion: we observed a mostly consistent downregulation across all time points, but this effect was not statistically significant in this context. In female embryos, however, we did observe a statistically significant decrease in *Xist* expression from the deletion alleles (**Fig. 6D-E**), and more pronounced for the Chic1-14kbΔ allele. This suggests that the *Chic1* CTCF cluster might operate to favour *Xist* expression in cis. These results potentially illustrate as well how loss of one additional CTCF binding site might be enough to cause stronger changes in chromosome conformation and gene expression.

**Figure 6.**
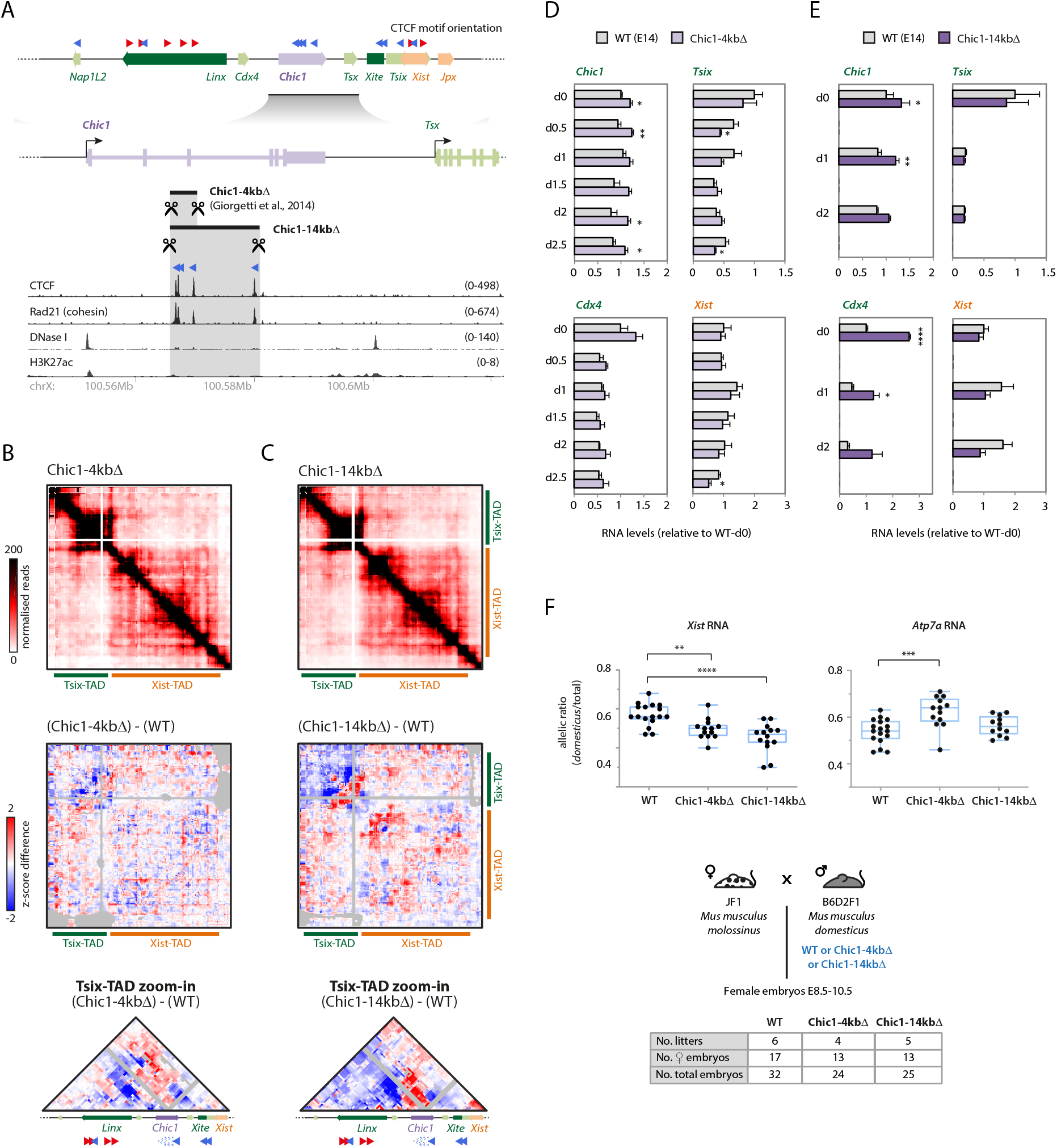
Deletion of *Chic1* cluster of CTCF sites leads to Xist downregulation in *cis*. (*A*) The *Chic1* locus, CTCF binding, and orientation of CTCF motifs associated with CTCF chromatin immunoprecipitation sequencing (ChIP-seq) peaks. The targeted deletions Chic1-4kbΔ and Chic1-14kbΔ are indicated. (*B, C*) Top, 5C profiles of Chic1-4kbΔ (*B*, two clones pooled) and Chic1-14kbΔ (*C*, one clone, two replicates pooled). Middle, 5C differential maps, representing the subtraction of Z-scores calculated for wildtype and deletion maps. Grey pixels represent filtered contacts (see Methods). Bottom, Tsix-TAD zoom-in of differential maps. (*D, E*) Gene expression analysis during differentiation. Data is normalised to wt-d0 for each gene, and represents the average of two biological replicates (wild type and Chic1-14kbΔ) or of two independent clones (Chic1-4kbΔ). Statistical analysis: two-tailed paired t-test (* p<0.05; ** p<0.01; **** p<0.0001). (*F*) Top, RNA allelic ratios for *Xist* and the X-linked gene *Atp7a*. Each black dot corresponds to a single female embryo. Statistical analysis was performed using Mann-Whitney test (** p<0.01; *** p<0.001; **** p<0.0001). Bottom, crosses used for analysis of RNA allelic ratios in female hybrid embryos inheriting the *M. musculus domesticus* allele paternally. Tables summarise number of embryos collected.

Together, our results on inverting or deleting *Linx* and *Chic1* CTCF clusters highlight the rather complex regulatory landscape within the *Xic*. Similar to the 245kb inversion, these mutant alleles reveal how *Xist* is sensitive to changes involving CTCF binding sites within the neighbouring Tsix-TAD. These results also suggest that the phenotypes observed in the 245kb-INV mESC are likely a compound of effects from changing different elements within the Tsix-TAD.

## Discussion

Here we explored the structural and transcriptional consequences of inverting a large genomic region encompassing almost an entire TAD (80%; 245kb out of 300kb). We found that this inversion led to rearrangement of contacts and to changes in expression of some genes within the TAD. We also observed increased contact insulation with the neighbouring TAD and ectopic upregulation of a gene in that TAD, the noncoding RNA *Xist* locus.

The rearrangement of contacts within the Tsix-TAD upon inverting a 245kb region occurred largely as expected based on the “rules” associated with the orientation of CTCF motifs within the TAD (de Wit et al., 2015; Guo et al., 2015; Rao et al., 2014a; Sanborn et al., 2015; Tang et al., 2015b). We found that the three loci involved in the strong contacts observed in the wild type Tsix-TAD were still able to form strong contacts with each other in the “inverted” Tsix-TAD (**Fig. 2**). Yet, these elements could not fully replace each other in their new positions, despite similar composition in terms of number of CTCF sites and levels of CTCF binding based on published ChIP-seq data (**Fig. 1**). In particular, the *Linx* CTCF cluster seems to have a stronger potential to form contacts than the *Chic1* CTCF cluster. At the same relative position within the TAD, and with the same CTCF motif orientation, these CTCF clusters show a different range of interactions, as described above (**Fig. 2**). These differences suggest that not all CTCF-bound sites are equally capable of mediating the same type of interactions. Little is known about what determines which CTCF sites contact with each other, and whether there could be specific affinities between sites – depending for instance on which other protein complexes are bound at each site or nearby. The ChIP-seq signal for CTCF is fairly comparable between CTCF sites within *Linx* and *Chic1*, and there are the same number of CTCF sites within each locus. One difference in the organisation of these sites is the spacing between them: CTCF sites within *Chic1* are more clustered than the ones within *Linx* – this could potentially play a role in orchestrating which and how contacts are formed. “Loop extrusion” has been proposed as a mechanism to form TADs and chromatin contacts (Fudenberg et al., 2016; Goloborodko et al., 2016; Sanborn et al., 2015), by which an “extruding factor” (such as cohesin) engulfs two DNA chains and moves along them, extruding DNA until it reaches “stalling factors” (such as CTCF), which block its progression; a chromatin loop would thus be formed and stabilized. Could the length of the intervals between CTCF sites influence the likelihood at which the cohesin complex (or the extruding factor) gets stalled? Perhaps more distributed sites (like at the *Linx* locus) rather than more clustered (like at the *Chic1* locus) provide more opportunities for stalling cohesin, given the very fast rate at which CTCF binds and unbinds chromatin (Hansen et al., 2017) and the rate of extrusion by cohesin (Davidson et al., 2019; Kim et al., 2019). Another potential explanation (not mutually exclusive) is that the differences in “contact potential” depend on the different sequences flanking the consensus CTCF motifs within *Linx* and *Chic1*, as suggested by a recent study on CTCF sites as transcriptional insulators (Huang et al., 2021).

Transcriptional changes in mutant 245kb-INV mESCs and during differentiation, when compared to control, were observed for two genes (*Nap1L2* and *Tsx*) within the Tsix-TAD (**Fig. 3**). As discussed previously, we think that these changes are associated with (genomic) proximity to the enhancer element *Xite* and not necessarily with the new topological structure of the inverted Tsix-TAD. Perhaps more interesting is the fact that most other genes within the Tsix-TAD do not show changes in expression, especially the *Linx, Chic1* and *Xite* loci, which are involved in the topological changes observed for the 245kb-INV allele. This could have a number of explanations: (i) the expression of these genes might not be particularly reliant on cis-regulation, and therefore impervious to topological changes; (ii) interactions between these genes and their cis-regulatory elements might not depend on topological organisation, and therefore still occur regardless of the topological changes; or (iii) interactions between these genes and their cis-regulatory elements might depend on topological organisation, and the new contacts allow these interactions to occur as efficiently as in wild type, and therefore no changes in expression are observed. Further genetic exploration of these loci will be critical to exclude hypotheses.

Surprisingly, expression of *Xist*, which lies outside of the Tsix-TAD, in the neighbouring TAD, was mildly upregulated, to an extent that we could detect accumulation of Xist RNA in “clouds” in mutant male cells, which we never observe(d) in wild type male cells (**Fig. 3**). This upregulation could be associated with one or more of the other alterations observed on the 245kb-INV allele, either structural, or transcriptional, or both. For instance, we observed reduced expression for *Tsix, Xist*’s antisense cis-repressor, in the pluripotent state, which could have an impact in *Xist* regulation; during differentiation, however, when *Xist* is upregulated, we did not detect differences in *Tsix* expression. Further research will be needed to clarify the involvement of *Tsix* in the *Xist* phenotype observed here.

Could *Xist* upregulation be a consequence of *Nap1L2* upregulation or *Tsx* downregulation? Genetic studies with *Nap1L2* (Attia et al., 2007; Rogner et al., 2000) did not report any effects on *Xist* expression or sex-specific phenotypes; upregulation of *Nap1L2* is thus unlikely to cause *Xist* upregulation, although this cannot be formally excluded. On the other hand, knockout studies of *Tsx* (Anguera et al., 2011) reported Xist RNA clouds in a small percentage of differentiating male mutant mESCs; the authors proposed that *Xist* expression was upregulated due to its negative cis-regulators *Tsix* and *Xite* being downregulated during differentiation. This *Xist* phenotype is identical to the one we observed (**Fig. 3**), although in 245kb-INV mutant cells there is still some *Tsx* expression (contrary to the *Tsx* knockout) and we did not observe changes in *Tsix* or *Xite* expression during differentiation. Downregulation of *Tsx* in 245kb-INV mutant cells might thus account, partially or maybe even completely, to ectopic *Xist* upregulation. This raises interesting questions of how such inter-TAD regulation/communication between *Tsx* and *Xist* could occur. Similarly, we have recently reported that another locus within the Tsix-TAD, *Linx*, contains sequences that affect expression of *Xist* in the neighbouring TAD in a *Tsix*-independent manner (Galupa et al., 2020). A slight increase in *Xist* expression in *cis* was also observed in 245kb-INV heterozygous embryos, but it was not statistically significant and did not result in skewed patterns of X-inactivation (**Fig. 4**). These results underlie the importance of verifying whether changes in gene expression result or not in differences of the phenotypes they mediate – in many studies it often remains an open question whether the changes observed in gene expression, especially when modest, do matter for the processes in which those genes are involved.

In agreement with previous studies, our study illustrates that the relationship between chromosome structure and gene expression is rather complex. The almost “dogmatic” view that TADs restrict gene cis-regulation (Finn and Misteli, 2019; Koch, 2019) is at odds with a growing amount of evidence that mechanisms of inter-TAD communication exist, albeit potentially subject to modulation by TADs and their boundaries. Here, we showed that, on the one hand, expression of genes within a TAD can be quite tolerant to changes in contacts within that TAD; on the other hand, we also showed that inversion of a large region within a TAD affected the expression of a gene in the neighbouring TAD, potentially due to accompanying changes in topological organisation and topological insulation. Further investigations are warranted for a more complete understanding of the relationship between the topological organisation of the genome and the transcriptional regulation of its genes.

## Materials and methods

All the materials and methods described below have also been published previously (Galupa et al., 2020).

### Tissue culture conditions

E14 mESC line and clones derived from it were grown on flasks or dishes coated with 0.1% (wt/vol) gelatin. Culture media for mESC consisted in Glasgow medium (Gibco) supplemented with 2mM L-Glutamine, 0.1mM nonessential amino acids, 1mM sodium pyruvate, 15% FBS (Gibco), 0.1 mM b-mercaptoethanol (Sigma) and 1000 U/mL of LIF (Chemicon). All lines were cultivated at 37ºC under 8% CO_2_ and passaged according to their confluency, generally every other day. Medium was refreshed daily. For early differentiation assays, mESC were washed with 1x PBS, incubated with trypsin at 37ºC for 20min and resuspended in ES medium without LIF. After cell counting, desired number of cells was resuspended in differentiation medium and 8*10^5^ cells per well were seeded in a fibronectin-coated (10 □g/mL, Millipore) 6-well plate in differentiation medium. Differentiation medium was consisted of N2B27 medium, 20 ng/mL activin A (R&D) and 12 ng/mL FGF-basic (R&D). Differentiation medium was changed daily and cells were washed in PBS before collection to remove dead cells.

### Mouse experimentation

Animal care and use for this study were performed in accordance with the recommendations of the European Community (2010/63/UE) for the care and use of laboratory animals. Experimental procedures, including genomic engineering (see below), are in compliance with international guidelines and were specifically approved by the ethics committee of the Institut Curie CEEA-IC #118 and given authorization by the French national authorities (references: APAFIS##13962-2018030717538778-v2 and APAFIS#8812-2017020611033784-v2).

Postimplantation embryos were collected at E8.5-10.5 stages, assuming plugging at midnight. Females with a vaginal plug were weighted every other day and only taken for dissection if a significant increase in weight was observed (∼2g for B6D2F1 mice, ∼1g for JF1 mice) at expected time of E8.5-E10.5 development. Extraembryonic tissues were taken for sexing the embryos. Whole embryo proper was washed three times in 1xPBS before frozen for allelic expression analysis.

### Genomic engineering of mice and mESC

Inversion 245kb-INV and deletion Chic1-14kbΔ were generated using CRISPR-Cas9 (mESC and mice) technologies, and the process is described below. Inversions within the *Linx* locus (Linx-25kb-INV and Linx-51kb-INV) were generated using the same constructs and primers as the equivalent deletions, which were described in (Galupa et al., 2020). The deletion Chic1-4kbΔ was previously generated (Giorgetti et al., 2014).

We designed sgRNAs to flank the region of interest:

- For 245kb-INV: CR30 (ACTGGTTCAGCCACTCACCG) and CR32 (CTGAGCTGGTTCATACAGGT)
- For Chic1-14kbΔ: CR21 (AAAGATCGTTTCTATCTAGC) and CR16R (CGCCAAACTTCCAAAATGGC)

For cloning sgRNAs, we used pX459-v2 (Plasmid #62988; Addgene) and protocol from the Zhang lab (https://media.addgene.org/cms/filer_public/e6/5a/e65a9ef8-c8ac-4f88-98da-3b7d7960394c/zhang-lab-general-cloning-protocol.pdf). sgRNA constructs were amplified upon transformation of DH5α competent cells (Takara) grown at 37ºC, and sequenced for verifying correct cloning. Midipreps for all constructs were prepared at final concentration >1mg/mL using the NucleoBond Xtra Midi Plus kit (Macherey-Nagel).

mESC were transfected with sgRNA constructs using the P3 Primary Cell 4D-Nucleofector X Kit (V4XP-3024) and the Amaxa 4D Nucleofector™ system (Lonza). We used the transfection programme CG-104. Each transfection included 5 million cells resuspended in the nucleofection mix (prepared according to manufacturer’s instructions) containing 5μg of each sgRNA (two constructs). As a transfection control, 10μg of pmaxGFP (Lonza) were used, for which the nucleofection efficiency was around 90%. Cells were immediately resuspended in pre-warmed culture medium after nucleofection and seeded at three serial 10x dilutions in 10-cm dishes to ensure optimal density for colony-picking. Transfected cells were selected with puromycin for 48h, and grown for 8-10 days. Single colonies were picked into 96-well plates. Genomic DNA was isolated in 96-well plates for PCR-based screening of inversions. Genotyping primers:

- For 245kb-INV: RG82 (CAATCACTCTTGCCTTACCAATT), RG83 (CCCAAACCAACCCTTGACTG), RG84 (GTTGGGACCTAAACTCTAGTACA) and RG85 (AGTGGACTAGCTTTGCCTCA)
- For Chic1-14kbΔ: EN118 (GCCTGCAGTCTTACCAGGAG), EN119 (TAATCTGCAGCGTGTTGAGG), RG123 (TCCTCCCTTACCAGTCTCCT), RG124 (CAGAATCCCGGATGTGAGGA)

The strategy was inspired on the Epigenesys protocol by Nora and Heard, 2012, described in: https://www.epigenesys.eu/en/protocols/genome-engineering/816-engineering-genomic-deletions-and-inversions-in-mouse-es-cells-using-custom-designed-nucleases.

We sequenced the PCR products from the inversion alleles to determine the breakpoints:

- For 245kb-INV – clone1: chrX-100377328 and chrX-100622017; clone2: chrX-100377337 and chrX-100622025 (coordinates in mm9)
- For Chic1-14kbΔ – clone1: chrX-103370850 and chrX-103384956 (coordinates mm10)

The mouse mutant lines were generated following the strategy described in (Wang et al., 2013) with minor modifications. Cas9 mRNA was in vitro transcribed from a T7-Cas9 pCR2.1-XL plasmid (Greenberg et al., 2017) using the mMESSAGE mMACHINE T7 ULTRA kit (Life Technologies) and purified with the RNeasy Mini kit (Qiagen), or bought from Tebu-bio (L-7206). The sgRNAs were amplified by PCR with primers containing a 5′ T7 promoter sequence from the plasmids used for mESC transfection. After gel purification, the T7-sgRNA PCR products were used as the template for in vitro transcription with the MEGAshortscript T7 kit (Life Technologies) and the products were purified using the MEGAclear kit (Life Technologies). Cas9 mRNA and the sgRNAs were eluted in DEPC-treated RNase-free water, and their quality was assessed by electrophoresis on an agarose gel after incubation at 95ºC for 3min with denaturing agent provided with the in vitro transcription kits. Cas9 mRNA and sgRNAs (at 100 ng/μl and 50 ng/μl, respectively) were injected into the cytoplasm of mouse B6D2F1 zygotes from eight-week-old superovulated B6D2F1 (C57BL/6J × DBA2) females mated to stud males of the same background. Zygotes with well-recognized pronuclei were collected in M2 medium (Sigma) at E0.5. Injected embryos were cultured in M16 medium (Sigma) at 37°C under 5% CO2, until transfer at the one-cell stage the same day or at the two-cell stage the following day to the infudibulum of the oviduct of a pseudogestant CD1 female at E0.5 (25-30 embryos were transferred per female). All weaned mice (N0) were genotyped for presence of inversion alleles using the same genotyping primers as for mESC mutant lines. Mice carrying inversion alleles were crossed to B6D2F1 mice and their progeny screened again for the presence of the inversion allele. The F1 mice were considered the “founders” and bred to B6D2F1 mice; their progeny was then intercrossed to generate homozygous mice and lines were kept in homozygosity.

### RNA fluorescent in situ hybridisation (FISH)

RNA FISH was performed as described previously with minor modifications (Chaumeil et al., 2008). Briefly, differentiating mESCs were dissociated using accutase (Invitrogen) and adsorbed onto Poly-L-Lysine (Sigma) coated coverslips #1.5 (1mm) for 5 min. Cells were fixed with 3% paraformaldehyde in PBS for 10 min at room temperature and permeabilized for 5 min on ice in PBS containing 0.5%Triton X-100 and 2mM Vanadylribonucleoside complex (New England Biolabs). Coverslips were preserved in 70% EtOH at -20°C. To start FISH experiments, coverslips were dehydrated through an ethanol series (80%, 95%, and 100% twice) and air-dried quickly, then lowered onto a drop of the probe/hybridization buffer mix (50% Formamide, 20% Dextran sulfate, 2x SSC, 1μg/μl BSA, 10mM Vanadyl-ribonucleoside) and incubated overnight at 37°C. The next day, coverslips were washed three times at 42 °C in 50% formamide in 2× SSC (pH 7.2-7.4) and three times at 42 °C in 2× SSC. Nuclei were counterstained with DAPI (0.2mg/ml), coverslips were mounted (90% glycerol, 0.1X PBS, 0.1% p-phenylenediamine at pH9), and cells were imaged using a wide-field DeltaVision Core microscope (Applied Precision).

Probes used were a *Huwe1* bacterial artificial chromosome, BAC (BACPAC Resources Center, RP24-157H12) and oligos (∼75 nucleotides long) covering all *Xist* exons (Roche, custom design). The BAC was labelled using the Nick Translation kit from Abbot and following manufacturer’s instructions. Oligos were end-labeled with Alexa488 fluorophore (from manufacturer). Probes were either ethanol-precipitated (BAC) or vacuum-dried (oligos) and resuspended in formamide with shaking at 37°C. BAC was co-precipitated with mouse Cot-1 DNA (Invitrogen), and competition to block repetitive sequences was performed for at least 20min at 37°C, and after denaturation (75°C, 10 min). Probes were then mixed with one volume of 2× hybridization buffer.

### Gene expression analysis (mESCs)

Cells were collected for gene expression analysis at 0h, 12h, 24h, 36h, 48h and 60h of differentiation. Cells were lysed with Trizol (Invitrogen), and RNA was isolated using the RNAeasy Mini kit (Qiagen), including DNase treatment. RNA samples were systematically run on an agarose gel to check their integrity. For reverse transcription cDNA was synthesised from 0.5μg of RNA using SuperScript™ III Reverse Transcriptase and random primers (both Invitrogen) according to the manufacturer’s recommendations. Two independent reverse transcription experiments were carried out for each sample, pooled at the end and diluted 25-fold prior to qPCR or allelic expression analysis. No-reverse transcription controls were processed in parallel. We used the NanoString nCounter gene expression system (Geiss et al., 2008) to systematically characterise transcriptional differences in wildtype and mutant mESC, prior or during differentiation. We used 500ng of total RNA from each sample for each nCounter hybridization round. We designed a customised probe codeset (van Bemmel et al., 2019) to identify nearly a hundred transcripts from *Xic* genes, other X-linked genes, pluripotency factors, differentiation markers, proliferation markers and normalization genes. Standard positive controls included in the kit were used for scaling the raw data. Genes Actb, Rrm2 and Sdha were used for normalization. Differential expression was always calculated for samples run on the same nCounter hybridization.

### Allelic expression analysis (mouse embryos)

Embryos were lysed in RLT buffer (Qiagen) supplemented with 0.01% 2-mercaptoethanol, and after two rounds of vortexing (15sec each), lysates were applied directly to a QIAshredder spin column (Qiagen) and centrifuged for 3min at full speed. RNA was extracted using the RNAeasy Mini kit (Qiagen), including DNase treatment, and following manufacturer’s instructions. RNA samples were systematically run on an agarose gel to check their integrity. cDNA was prepared as described above for gene expression analysis for mESCs, and then PCR-amplified with biotinylated primers and pyrosequenced for allele quantification on a Pyromark Q24 system (Qiagen). The same PCR was done on no-reverse transcription control samples to confirm absence of genomic DNA contamination. Primers used were designed using the PyroMark Assay Design software and validated on XX polymorphic genomic DNA for a ratio of 50:50% (± 4%). List of primers and SNPs used for allele quantification can be found in (Galupa et al., 2020).

### Chromosome conformation capture

3C libraries were prepared based on previous protocols (Nora et al., 2017; Rao et al., 2014b), with some modifications. Crosslinked cells (in 2% Formaldehyde; 10 million for each sample) were lysed in 10 mM Tris–HCl, pH 8, 10 mM NaCl, 0.2% NP-40, 1 × complete protease inhibitor cocktail (Roche) for 15min on ice. Nuclei were resuspended in 100 μL 0.5% SDS, incubated at 62°C for 10min and quenched with 50 μL 10% Triton X-100 and 290 μL water at 37°C for 15min. Digestion was performed overnight by adding 50 μL of DpnII (Capture-C) or HindIII (5C) buffer and 10 μL of high-concentration DpnII or HindIII (NEB) and incubating samples at 37°C in a thermomixer. Before this step, an aliquot was taken from each sample as an undigested control. Digests were heat inactivated for 20 min at 65ºC and an aliquot was taken from each sample as a digested (unligated) control. Samples were cooled at room temperature for 10 min before adding the ligation cocktail. 3C libraries were ligated for 4 hours at 25ºC with 10U T4 ligase and ligation buffer (ThermoFisher cat 15224) in a thermomixer at 1000rpm. Ligated samples were then centrifuged at 2000rpm, resuspended in 240 μL of 5% SDS and 1 mg Proteinase K, incubated at 55ºC for 30min, supplemented with 50 μL 5 M NaCl and incubated at 65ºC for 4 hours. DNA was then purified by adding 500 μL isopropanol, incubated at -80ºC overnight, centrifuged at 12,000 rpm at 4ºC, washed with 70% ethanol, air dried and resuspended in 100 μL water, followed by incubation with RNase A at 37ºC for one hour. 3C templates were quantified using Qubit DNA Broad-Range (ThermoFisher) and diluted to 100 ng/μL. Libraries and respective controls (undigested and digested aliquots) were verified on a gel.

5C was performed as described in (Nora et al., 2017), which adopts a single-PCR strategy to construct 5C-sequencing libraries from the 3C template. Briefly, four 10 μL 5C annealing reactions were assembled in parallel, each using 500 ng of 3C template, 1 μg salmon sperm (ThermoFisher) and 10 fmol of each 5C oligonucleotide in 1X NEBuffer™ 4 (5C set of oligonucleotides described in Nora et al., 2012). Samples were denatured at 95ºC for 5 min and incubated at 48ºC for 16-18h. 10 μL of 1X Taq ligase buffer with 5U Taq ligase were added to each annealing reaction followed by incubation at 48ºC for 4h and 65ºC for 10 min. Negative controls (no ligase, no template or no 5C oligonucleotide) were included during each experiment to ensure the absence of contamination. To attach Illumina-compatible sequences, 5C libraries were directly PCR amplified with primers harboring 50-mer tails containing Illumina sequences that anneal to the universal T3/T7 portion of the 5C oligonucleotides (Nora et al., 2017). For this, each 5C ligation reaction was used as the template for three parallel PCRs (12 PCRs total), using per reaction 6 μL of 5C ligation with 1.125 U AmpliTaq Gold (ThermoFisher) in 1X PCR buffer II, 1.8 mM MgCl2, 0.2 mM dNTPs, 1.25 mM primers in 25 mL total. Cycling conditions were 95ºC for 9 min, 25 cycles of 95ºC for 30 sec, 60ºC for 30 sec, 72ºC for 30 sec followed by 72ºC for 8 min. PCR products from the same 3C sample were pooled and run on a 2.0% agarose electrophoresis gel. 5C libraries (231 bp) were then excised and purified with the MinElute Gel Extraction kit (QIAGEN). Library concentrations were estimated using TapeStation (Agilent) and Qubit (ThermoFisher), pooled and sequenced using 12 pM for the loading on rapid flow cells using the HiSeq 2500 system (Illumina). Sequencing mode was set as 20 dark cycles followed by 80 bases in single end reads (SR80).

Sequencing data was processed using our custom pipeline, 5C-Pro, available at https://github.com/bioinfo-pf-curie/5C-Pro. Briefly, single-end sequencing reads were first trimmed to remove Illumina adapters and aligned on an *in silico* reference of all pairs of forward and reverse primers using the bowtie2 software (Langmead and Salzberg, 2012). Aligned reads were then directly used to infer the number of contacts between pairs of forward and reverse primers, thus providing a 5C map at the primer resolution. Based on our previous experiments, inefficient primers were discarded from downstream analysis. Quality controls of the experiments were then performed using the HiTC BioConductor package (Servant et al., 2012). Data from biological replicates were pooled (summed) and binned using a running median (window=30kb, final resolution=6kb). We normalized 5C contacts for the total number of reads and filtered out outlier probes and singletons, as previously described (Hnisz et al., 2016; Nora et al., 2012; Smith et al., 2016). We also developed a novel method to exclude noisy contacts in the 5C maps, called “neighbourhood coefficient of variation”, available at https://github.com/zhanyinx/Coefficient_Variation. Considering that the chromatin fiber behaves as a polymer, the contact frequency of a given pair of genomic loci (e.g. *i* and *j*) cannot be very different from those of fragments *i*±*N* and *j*±*N* if *N* is smaller (or in the order of) than the persistence length of the chromatin fiber. Hence, a given pixel in the 5C map (which is proportional to the contact frequency between the two corresponding loci) can be defined as noisy if its numerical value is too different from those corresponding to neighboring interaction frequencies. To operatively assess the similarity of a given interaction with neighboring contacts, we calculated the coefficient of variation (CV) of contacts (pixels in the 5C map) in a 10×10 square centered on every contact. We then set out to discard pixels for which the corresponding coefficient of variation was bigger than a threshold. Given that the distribution of the coefficient of variation of all 5C samples in this study is bimodal around CV=1, we set the CV threshold to 1. Discarded contacts appear as grey pixels in the differential 5C maps. For differential analysis between two samples of interest, we calculated the difference between Z-scores determined for each individual map (Smith et al., 2016). Samples corresponding to inversions of genomic regions were mapped to a virtually inverted map before analysis. Samples corresponding to deletions were corrected for the new distance between genomic elements; this distance-adjustment was performed along with the Z-score calculation. 5C data for E14 cell line (used as control) has been published previously (Galupa et al., 2020) but control and mutant samples were collected and processed in parallel.

### Statistical analysis

For RNA FISH, nCounter and allelic expression analysis, statistical details of experiments can be found in the figure legends, figures and/or Results, including the statistical tests used, exact value of n and what n represents.

### Accession numbers

All next-generation sequencing data generated in this study has been deposited in the Gene Expression Omnibus (GEO) under the accession number GSE124596 (5C data for E14 cell line) and GSE180617 (5C data for 245kb-INV cell line). 5C data for E14 cell line (used as control) has been published previously (Galupa et al., 2020) but control and mutant samples were collected and processed in parallel.

----------

To review GEO accession GSE180617:

Go to https://www.ncbi.nlm.nih.gov/geo/query/acc.cgi?acc=GSE180617 Enter token gdelimmatputvod into the box

----------

## Acknowledgments

We are grateful to Isabelle Grandjean for help and advice with animal management; to Patricia Diabangouaya for help with mouse genotyping; to Chris Gard for help with gene expression analysis; to Denis Krndija for critical reading of the manuscript. We thank all members of the Heard lab for advice, support, and helpful comments and discussions. We are also thankful to facilities at the Institut Curie, including the Mouse Facility, the BDD team of PICT-IBiSA, the ICGex NGS platform (in particular Sónia Lameiras and Sylvain Baulande), the Genomics Platform (in particular David Gentien, Cécile Reyes, Audrey Rapinat and Benoit Albaud) and the Bioinformatics Platform. We acknowledge the Zhang lab for sharing plasmids, and the ENCODE Consortium and the Bruneau, Ren, Sharp, Stamatoyannopoulos and Young labs for generating datasets used in this study.

## Author contributions

Conceptualization: RG, LG, EH. Investigation: RG, CP, EPN. Methodology: RG, CP, EPN, FEM, CJ, MB, KA. Formal analysis: RG, NS, YZ, JvB. Data curation: RG, NS, YZ, JvB, LG. Visualization: RG, YZ. Software: NS, YZ, JvB, LG. Supervision: RG, MB, KA, LG, EH. Project administration: RG, EH. Funding acquisition: EH. Writing, original draft: RG. Writing, review & editing: RG, KA, LG, EH.

## Figure legends

**Figure S1**. (*A*) Virtual 4C plots for the wild type (WT) and 245kb-INV alleles, for which the anchor is the element *Xite*. The interaction frequency between *Xite-Tsx* is ∼5-fold lower in the inverted allele (∼50 counts) than in the WT (∼250 counts, and within the region where contact frequency is dominated by genomic distance). Inversely, the interaction frequency between *Xite-Nap1L2* in the inverted allele (∼250 counts) is ∼5-fold higher than in the WT (∼50 counts). The changes in interaction frequencies between these elements seem thus to reflect the changes in genomic distances for WT and inverted alleles. (*A*) Virtual 4C plots for the wild type (WT) and deletion alleles Chic1-4kbD and Chic1-14kbD. The interaction frequency between *Xite* and *Linx* is increased in the mutant alleles compared to WT.

